# Homeostatic regulation of STING by Golgi-to-ER membrane traffic

**DOI:** 10.1101/2020.05.20.107664

**Authors:** Kojiro Mukai, Emari Ogawa, Rei Uematsu, Yoshihiko Kuchitsu, Takefumi Uemura, Satoshi Waguri, Takehiro Suzuki, Naoshi Dohmae, Hiroyuki Arai, Anthony K. Shum, Tomohiko Taguchi

**Affiliations:** Laboratory of Organelle Pathophysiology, Department of Integrative Life Sciences, Graduate School of Life Sciences, Tohoku University, Sendai, Japan; Department of Health Chemistry, Graduate School of Pharmaceutical Sciences, University of Tokyo, Tokyo, Japan; Department of Anatomy and Histology, Fukushima Medical University School of Medicine, Fukushima, Japan; Biomolecular Characterization Unit, RIKEN Center for Sustainable Resource Science, Wako, Japan; Department of Medicine, Division of Pulmonary and Critical Care, University of California San Francisco, San Francisco, CA, USA

## Abstract

Coat protein complex I (COP-I) mediates the retrograde transport from the Golgi to the ER^1,2^. Mutation of the *COPA* gene, encoding one of the COP-I subunits (α-COP), causes an immune dysregulatory disease (COPA syndrome)^3^. The molecular mechanism by which the impaired retrograde transport results in autoinflammation is not understood. Here we report that STING^4^, an innate immunity protein, is a cargo of the Golgi-to-ER membrane transport. In the presence of the disease-causative α-COP variants, STING cannot be retrieved back to the ER from the Golgi. The forced Golgi residency of STING results in the cGAS-independent and palmitoylation-dependent activation of the STING downstream signalling pathway. Surf4^5^, a protein that circulates between the ER and the Golgi, binds STING and α-COP, and mediates retrograde transport of STING to the ER. STING/Surf4/α-COP complex is disrupted in the presence of the disease-causative α-COP variant. Intriguingly, the STING ligand cGAMP also impairs the formation of STING/Surf4/α-COP complex. Our results suggest a homeostatic regulation of STING at the resting state by the Golgi-to-ER membrane traffic and provide insights into the pathogenesis of COPA syndrome.

---

The COPA syndrome is a recently discovered monogenic disorder of immune dysregulation characterized by high-titer autoantibodies, interstitial lung disease, inflammatory arthritis, and high expression of type I interferon-stimulated genes^3,6^. The disease is caused by heterozygous mutations of the *COPA* gene, encoding the α subunit (α-COP) of COP-I that mediates the retrograde transport of proteins from the Golgi to the endoplasmic reticulum (ER)^1,2^. How the retrograde transport in the COPA syndrome causes the immune dysregulatory disease remains largely unknown.

Vertebrates have evolved biological systems to combat invading pathogens. As the first line of host defense, the innate immune system detects microbial pathogens with pattern recognition receptors (PRRs) that bind unique pathogen-associated molecular patterns (PAMPs)^7,8^. Activated PRRs initiates intracellular signalling cascades, leading to the transcriptional expression of proinflammatory cytokines, type I interferons, and other antiviral proteins that all coordinate the elimination of pathogens and infected cells. Viral RNA, cytosolic DNA, or the gram-negative bacterial cell-wall component lipopolysaccharide serves as PAMP that activates a distinct signalling pathway, such as RIG-I/MAVS, cGAS/STING, or TLR4/TRIF pathway. MAVS, STING, or TRIF activates the downstream protein kinase TBK1, which then phosphorylates and activates interferon regulatory factor 3 (IRF3), the essential transcription factor that drives type I interferon production^9^.

STING^4^ is an ER-localized transmembrane protein. After STING binding to cyclic dinucleotides (CDNs)^10^ that are generated by cGAMP synthase (cGAS)^11^, an enzyme that is activated by the presence of cytosolic DNA, STING translocates to the Golgi where STING activates TBK1^12–14^. Because α-COP is a component of COP-I that mediates the membrane transport between the Golgi and the ER, we reasoned that the disease-causative α-COP variant (K230N, R233H, E241K, or D243G) (the α-COP variant hereafter)^3^ could influence the STING pathway.

We performed luciferase assay with HEK293T cells that lack endogenous STING. After co-transfection with α-COP, STING, and a luciferase reporter construct with IRF3 (also known as ISRE or PRD III-I)-responsive promoter elements, the luciferase activity in the total cell lysate was measured. Wild-type α-COP did not activate the IRF3 promoter regardless of the expression of STING, while all the α-COP variants activated the IRF3 promoter in STING-expressing cells (Fig. 1a). The α-COP variants did not activate the IRF3 promoter in cells transfected with MAVS or TRIF (Supplementary Fig. 1). We then examined if the expression of the α-COP variants results in TBK1 phosphorylation. When wild-type α-COP or the α-COP variants was stably expressed in *Sting*^-/-^ mouse embryonic fibroblasts (MEFs) or *Sting*^-/-^ MEFs reconstituted of EGFP-STING, phosphorylated TBK1 (p-TBK1) was observed only in cells expressing the α-COP variants and STING (Fig. 1b). These results suggested that expression of the α-COP variants could activate the STING pathway.

**Fig. 1.**
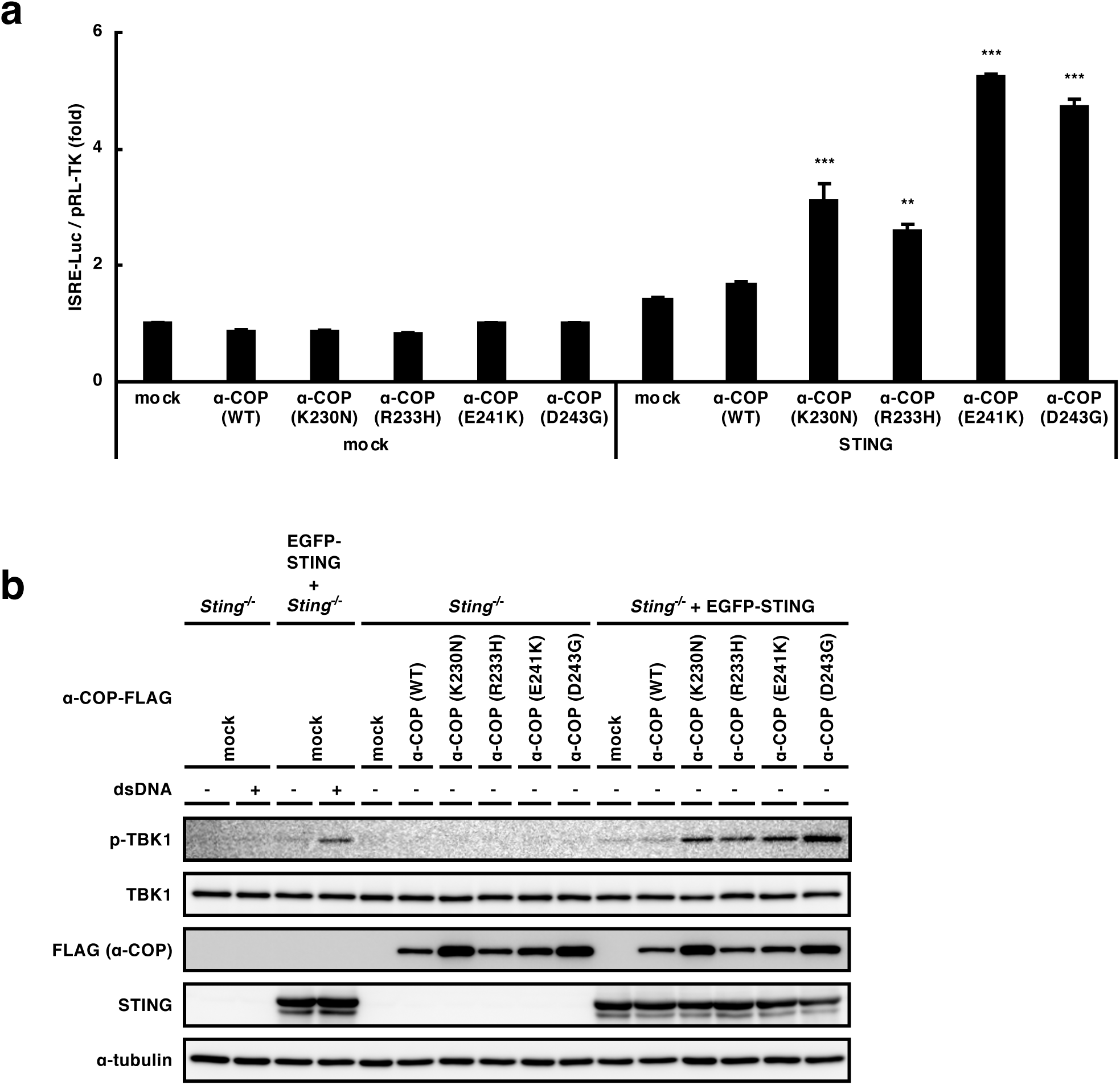
The α-COP variants activate the STING pathway. **a**, HEK293T cells were transfected as indicated, together with an ISRE (also known as PRDIII or IRF-E)-luciferase reporter. Luciferase activity was then measured. Data represent mean s.e.m. of three independent experiments. **b**, STING and/or α-COP were stably expressed in *Sting*^-/-^ MEF cells. Cell lysates were prepared and analyzed by western blot.

We next examined the subcellular localization of STING. In cells expressing wild-type α-COP, EGFP-STING distributed throughout the cytoplasm and co-localized with calreticulin (an ER protein), indicating that STING localized at the ER (Fig. 2a, Supplementary Fig. 2). p-TBK1 and phosphorylated STING at Ser365 (p-STING) that is generated by active TBK1^9,15^ were not detected. In contrast, in cells expressing the α-COP variants, EGFP-STING mostly localized at perinuclear compartments that include the Golgi (Fig. 2a, Supplementary Figs. 2-4). Thus, the expression of the α-COP variants altered the STING localization. The signals of p-TBK1 and p-STING emerged in these cells (Fig. 2b, Supplementary Figs. 5, 6), being consistent with the activation of STING (Fig. 1b). Immunoelectron microscopy corroborated the Golgi localization of STING in cells expressing the α-COP variant (E241K) (Fig. 2c). Given that COP-I mediates the retrograde transport from the Golgi to the ER^2^, these results suggested that STING is a cargo of the retrograde transport of COP-I and that STING cannot be retrieved back to the ER in the presence of the disease-causative α-COP variants. The amount of STING that was co-immunoprecipitated with the α-COP variants was smaller than that with wild-type α-COP (Fig. 2d), further supporting this notion.

**Fig. 2.**
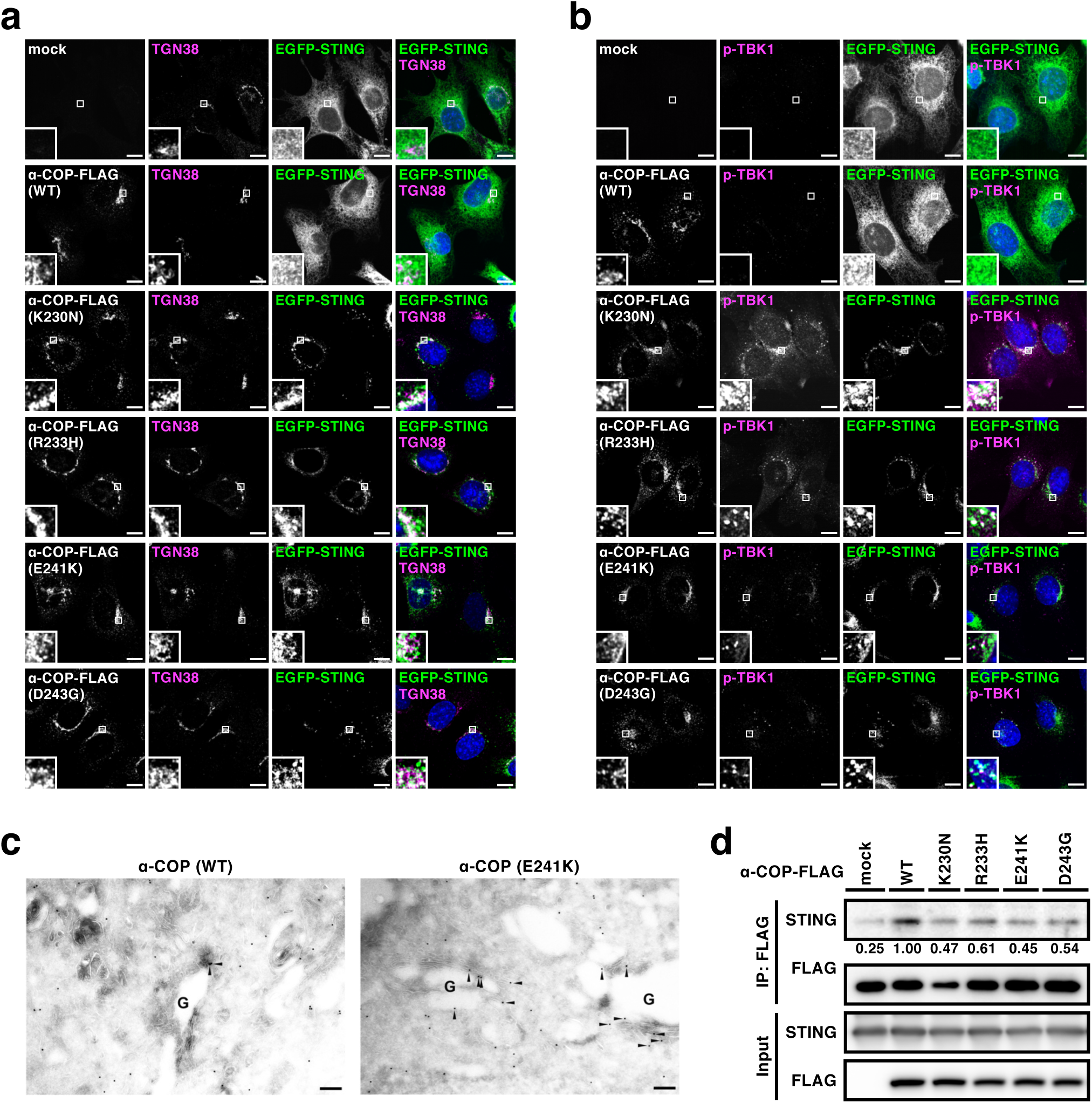
The α-COP variants alter STING localization to the Golgi. **a, b**, α-COP and EGFP-STING were stably expressed in *Sting*^-/-^ MEF cells as indicated. Cells were fixed, permeabilized, and stained for TGN38 (a Golgi protein) (**a**) or p-TBK1 (**b**). Nuclei were stained with DAPI (blue). Scale bars, 10 µm. **c**, α-COP (WT) or α-COP (E241K) were stably expressed in EGFP-STING-reconstituted MEF cells. Cells were fixed and processed for ultrathin-cryosections. They were immunostained with anti-GFP (rabbit) antibodies. As secondary antibodies, colloidal gold particle-conjugated donkey anti-rabbit antibody (12 nm) was used. Arrowheads indicate GFP labelling on the Golgi cisternae. G, the Golgi stack. Scale bars, 200 nm. **d**, Cell lysates were prepared from MEF cells expressing various α-COP as indicated, and α-COP was immunoprecipitated with anti-FLAG antibody. Cell lysates and the immunoprecipitates were analyzed by western blot.

α-COP binds *C*-terminal di-lysine motifs of its cargo proteins, such as KKXX and KXKXX^1,16,17^. As STING does not possess these motifs at its *C*-terminus, we reasoned the presence of adaptor protein(s) that mediates the interaction of STING and α-COP. We analyzed STING-binding proteins by mass spectrometry and 18 proteins with these motifs were identified (Fig. 3a). We knock-downed these proteins individually with siRNAs and examined the effect on the subcellular localization of STING. We found that knockdown of Surf4, not that of the other 17 proteins, altered the localization of STING to the Golgi (Fig. 3b, c, Supplementary Figs. 7, 8). Knockdown of Surf4 also resulted in the emergence of p-TBK1 in *Sting*^-/-^ MEFs reconstituted of EGFP-STING, but not in *Sting*^-/-^ MEFs (Fig. 3d). These results suggested that Surf4 is critical to maintain the steady-state localization of STING to the ER.

**Fig. 3.**
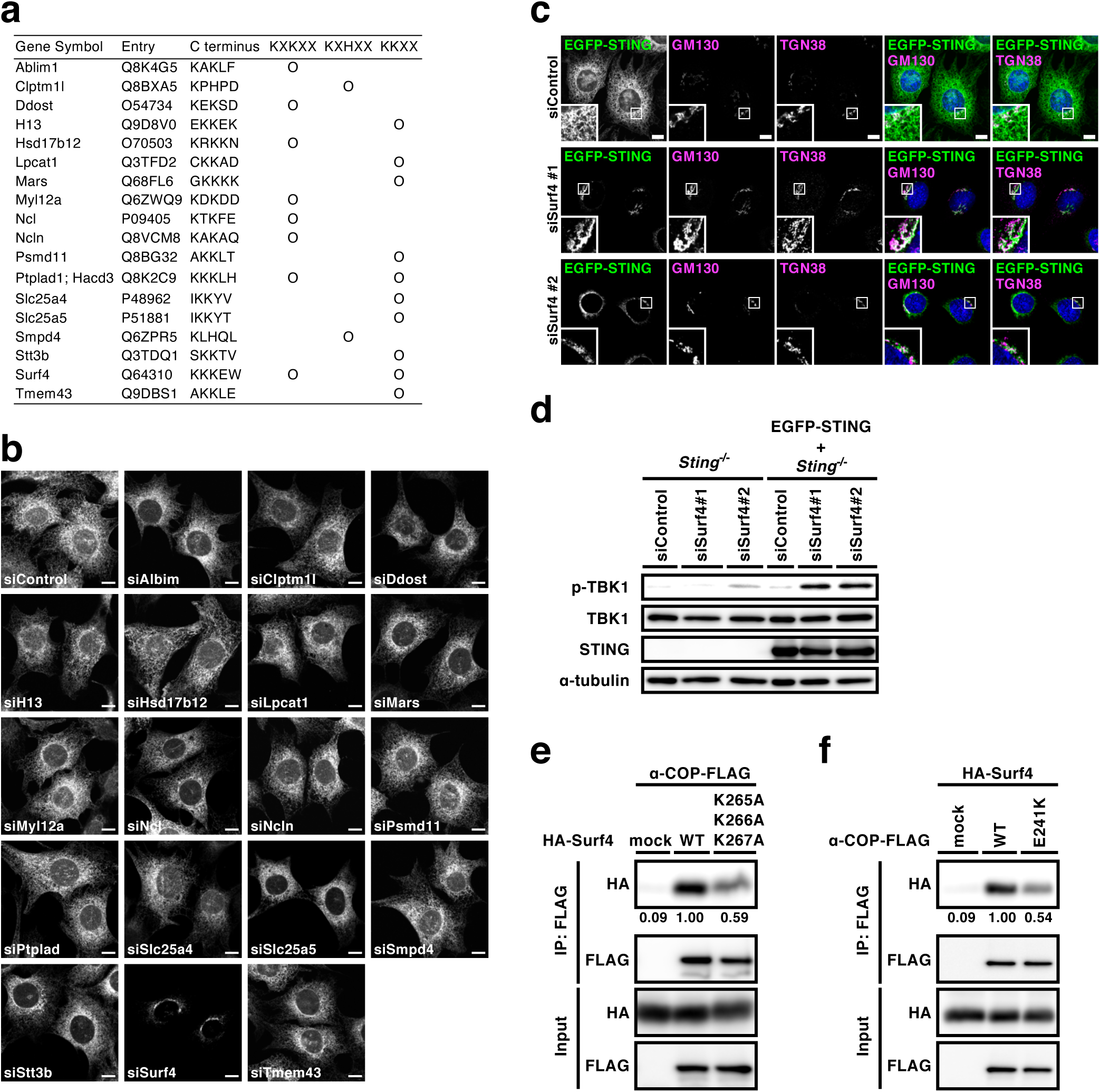
Surf4 binds STING and α-COP, and is required for STING localization to the ER. **a**, Cell lysates of FLAG-STING-reconstituted *Sting*^-/-^ MEF cells were prepared and FLAG-STING were immunoprecipitated with anti-FLAG M2 antibody. The immunoprecipitates were analyzed by mass spectrometry. Eighteen proteins with the COP-I binding motifs were identified and listed. **b, c**, MEF cells expressing EGFP-STING were treated with the indicated siRNA for 48 h. Cells were fixed, permeabilized, and stained for GM130 and TGN38. Scale bars, 10 µm. **d**, *Sting*^-/-^ MEF cells or *Sting*^-/-^ MEF cells reconstituted of EGFP-STING were treated with the indicated siRNA. Cell lysates were prepared and analyzed by western blotting. **e, f**, HEK293T cells were transfected with the indicated plasmids. Cell lysates were prepared and α-COP-FLAG was immunoprecipitated. Cell lysates and the immunoprecipitates were analyzed by western blot.

Surf4, a multi-pass transmembrane protein, cycles between the ER and the Golgi and is involved in membrane recruitment of COP-I^5^. Indeed, we found the interaction between Surf4 and wild-type α-COP by co-immunoprecipitation analysis (Fig. 3e). The mutant Surf4 (K265A/K266A/K267A), in which the *C*-terminal lysine residues were substituted to alanine, exhibited a reduced binding to α-COP, suggesting that the interaction between Surf4 and COPA is, at least in part, mediated through the *C*-terminal consecutive lysine residues on Surf4 (Fig. 3e). The disease-causative α-COP variant (E241K) exhibited a reduced binding to Surf4 (Fig. 3f). Given that Surf4 binds STING (Fig. 3a) and α-COP (Fig. 3e), Surf4 would serve as a cargo receptor for STING in the retrograde transport mediated by COP-I.

Activation of the STING signalling pathway with cGAMP requires the ER-to-Golgi traffic of STING and palmitoylation of STING at the Golgi^12,13,18^. We treated cells expressing the α-COP variants with brefeldin A (BFA), an agent to block the ER-to-Golgi traffic^19^, or two palmitoylation inhibitors [a pan-palmitoylation inhibitor 2-bromopalmitate (2-BP) and a mouse STING-specific palmitoylation inhibitor C-178^20^] and found that all the treatments suppressed phosphorylation of TBK1 (Fig. 4a, b, Supplementary Fig. 9). These results suggested that STING activation with the α-COP variants, as with cGAMP, requires the ER-to-Golgi traffic and palmitoylation of STING.

**Fig. 4.**
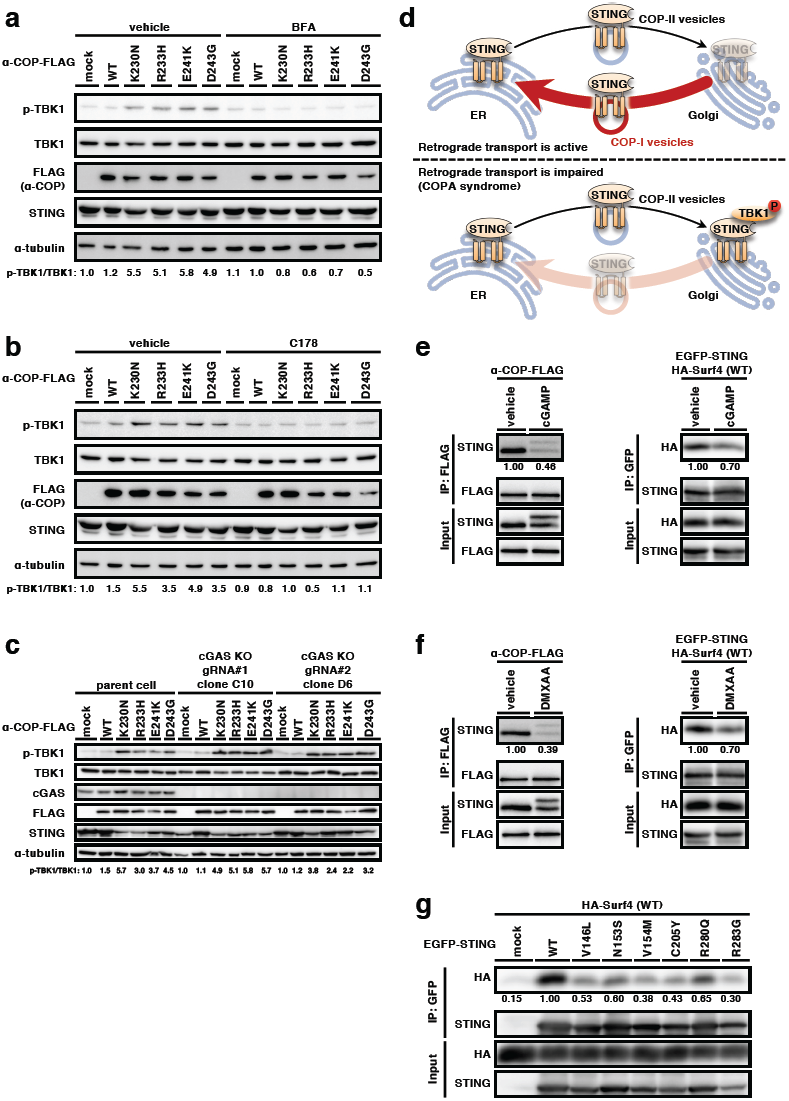
STING activation with the α-COP variants requires the ER-to-Golgi traffic and palmitoylation of STING, but not cGAS. **a, b**, MEF cells expressing various α-COP as indicated were treated with BFA (0.3 µg/ml) for 8 h (**a**) or with C178 (10 µM) for 8 h (**b**). Cell lysates were prepared and analyzed by western blotting. **c**, α-COP was stably expressed in cGAS-KO MEF cells. Cell lysates were prepared and analyzed by western blot. **d**, A model of STING regulation by membrane traffic between the ER and the Golgi. **e, f**, α-COP-FLAG or HA-Surf4 was stably expressed in *Sting*^-/-^ MEF cells reconstituted of EGFP-STING. Cells were stimulated with cGAMP (**e**) or DMXAA (**f**) for 1 h. Cell lysates were prepared, and α-COP-FLAG or EGFP-STING was immunoprecipitated. Cell lysates and the immunoprecipitates were analyzed by western blot. **g**, HEK293T cells were transfected with the indicated plasmids. Cell lysates were prepared and EGFP-STING were immunoprecipitated. Cell lysates and the immunoprecipitates were analyzed by western blot.

We asked if the STING activation with the α-COP variants requires cGAMP. To address this question, we prepared cGAS-knockout MEFs by CRISPR-Cas9 system. In cGAS-knockout MEFs expressing the α-COP variants, p-TBK1 still emerged (Fig. 4c) and STING translocated to perinuclear compartments that include the Golgi (Supplementary Fig. 10). These results suggested that the translocation of STING from the ER to the Golgi does not necessarily require cGAMP and that the forced Golgi residency of STING with the impaired retrograde transport would suffice to activate the STING signalling pathway in the absence of cGAMP.

These results led us to propose a model to explain how the membrane traffic axis between the ER and the Golgi is integrated into the STING signalling pathway (Fig. 4d). STING exits the ER without cGAMP. Once STING reaches the Golgi, STING is retrieved back to the ER by the COP-I-mediated retrograde transport. In a condition where the retrograde transport is impaired, such as in the presence of the disease-causative α-COP variants, STING is forced to accumulate at the Golgi. STING is then subjected to palmitoylation and activates TBK1.

Intriguingly, STING with DMXAA (a mouse STING agonist) or with cGAMP exhibited a reduced binding to α-COP and Surf4 (Fig. 4e, f). Therefore, the translocation of STING from the ER and activation of the STING signalling pathway with cGAMP may be partly due to the reduced ability of the STING/ligand complex to be packaged into COP-I transport vesicles. Mutations in STING are found in patients with an autoinflammatory disease called STING-associated vasculopathy with onset in infancy (SAVI) and these mutations appear to make STING constitutively active^21–23^. The SAVI variants exit the ER without cGAMP^13^ and require the ER-to-Golgi traffic and palmitoylation for their activity^12,13^. We found that all the six SAVI variants exhibited a reduced binding to Surf4 (Fig. 4g). The reduced binding may result in the impaired retrograde transport of STING to the ER, which would partly explain the aberrant localization of the SAVI variants to the Golgi without cGAMP.

In this study, we demonstrate the homeostatic regulation of STING at the resting state by Golgi-to-ER membrane traffic. Intriguingly, this finding is corroborated in a recently described mouse model of COPA syndrome (*Copa*^*E241K/*+^ mice)^24^: *Copa*^*E241K/*+^ mice exhibit spontaneous activation of STING with upregulation of type I interferon signalling and systemic inflammation, all of which is abrogated in STING-deficient animals^25^. Our results that the inflammatory response in the presence of COPA variants can be effectively suppressed by STING palmitoylation inhibitors may provide a new treatment approach for COPA syndrome patients^25^.

## Acknowledgements

This work was supported by JSPS KAKENHI Grant Numbers JP19H00974 (T.T.), JP15H05903 (T.T.), JP17H06164 (H.A.), JP17H06418 (H.A.), and JP17K15445 (K.M.); AMED-PRIME (17939604) (T.T.); Takeda Science Foundation (to S.W.). We thank Atsuko Yabashi for her technical support in the immuno-EM.

## Author contribution

K.M. designed and performed the experiments, analyzed the data, interpreted the results, and wrote the paper; E.O. designed and performed experiments, analyzed data, and interpreted results; R.U. performed the experiments for identification of STING binding proteins; Y.K. performed the experiments with cGAS KO cells; T.U. and S.W. performed the experiments with electron microscopy; T.S. and N.D. performed the proteomics analysis; H.A. designed the experiments, interpreted the results; A.K.S. discussed the results; T.T. designed the experiments, interpreted the results, and wrote the paper.

## Methods

### Antibodies

Antibodies used in this study were as follows: mouse anti-GFP (JL-8, dilution 1:1000) (Clontech); Alexa 488-, 594-, or 647-conjugated secondary antibodies (A21202, A21203, A21206, A21207, A31573, A11016, A21448, dilution 1:2000) (Thermo Fisher Scientific); rabbit anti-TBK1 (ab40676, dilution 1:1000) (Abcam); rabbit anti-phospho-TBK1 (D52C2, dilution 1:1000 for Western blotting, dilution 1:100 for immunofluorescence), rabbit anti-cGAS (D3O8O, dilution 1:1000), rabbit anti-phospho-STING (D1C4T, dilution 1:400) (Cell signaling); mouse anti-calreticulin (612136, dilution 1:1000), and mouse anti-GM130 (610823, dilution 1:1000) (BD Biosciences); mouse anti-α-tubulin (DM1A, dilution 1:5000) and mouse anti-FLAG M2 antibody (Sigma); Goat Anti-Rabbit IgG(H+L) Mouse/Human ads-HRP (4050-05, dilution 1:10000) and Goat Anti-Mouse IgG(H+L) Human ads-HRP (1031-05, dilution 1:10000) (Southern Biotech); sheep anti-TGN38 (AHR499G, dilution 1:500) (Serotec); rabbit anti-STING antibody (19851-1-AP, dilution 1:1000 for Western blotting) (Proteintech); mouse anti-HA (4B2, dilution 1:1000 for Western blotting and immunofluorescence) and mouse anti-FLAG (1E6, dilution 1:1000 for Western blotting and immunofluorescence) (Wako); colloidal gold particle-conjugated donkey anti-rabbit antibody (12 nm) (711-205-152, dilution 1:20) (Jackson ImmunoResearch laboratories). For the immunoprecipitation of FLAG-tagged protein, anti-FLAG M2 Affinity Gel (A2220, Sigma) was used. For the immunoprecipitation of GFP-tagged protein, anti-GFP nanobody was used. pGEX6P1-GFP-Nanobody was a gift from Kazuhisa Nakayama (Addgene plasmid # 61838).

### Reagents

The following reagents were purchased from the manufacturers as noted: BFA (Sigma); 2-BP (Wako). ISD (90-mer), used as dsDNA in this study, was prepared as follows: equimolar amounts of oligonucleotides (sense: 5’-TACAGATCTACTAGTGATCTATGACTGATCTGTACATGATCTACATACAGATCTACTAGTGATCTATGACTGATCTGTACATGATCTACA-3’, antisense: 5’-TGTAGATCATGTACAGATCAGTCATAGATCACTAGTAGATCTGTATGTAGATCATGTACAGATCAGTCATAGATCACTAGTAGATCTGTA-3’) were annealed in PBS at 70 °C for 30 min before cooling to room temperature. C-178 was provided by Carna Biosciences, Inc.

### Cell culture

HEK293T cells were purchased from the American Type Culture Collection (ATCC). MEFs were obtained from embryos of WT or *Sting*^*-/-*^ mice at E13.5 and immortalized with SV40 Large T antigen. HEK293T and MEFs were cultured in DMEM supplemented with 10% fetal bovine serum/penicillin/streptomycin/glutamine in a 5% CO_2_ incubator.

MEFs that stably express EGFP-mouse STING or mouse α-COP variants were established using retrovirus. Plat-E cells were transfected with pMX-IP-EGFP-STING or pMX-IB-α-COP-FLAG and the medium that contains the retrovirus was collected. MEFs were incubated with the medium and then selected with puromycin (2 µg/mL) or blasticidin (5 µg/mL) for several days.

### PCR cloning

Mouse STING was amplified by PCR with complementary DNA (cDNA) derived from ICR mouse liver. The product encoding mouse STING was introduced into pMXs-IPuro–GFP, to generate N-terminal GFP-tagged construct. Mouse α-COP and mouse Surf4 was amplified by polymerase chain reaction (PCR) with cDNA derived from MEFs. The product encoding α-COP was introduced into pMXs-IBla-FLAG, to generate C-terminal FLAG-tagged construct. The product encoding Surf4 was introduced into pMXs-IHyg-HA, to generate N-terminal HA-tagged construct. α-COP variants and Surf4 mutant were generated by site-directed mutagenesis.

### Luciferase assay

HEK293T cells seeded on 24-well plates were transiently transfected with luciferase reporter plasmid (100 ng), pRL-TK (10 ng) as internal control, STING-expression plasmid in pBabe vector (200 ng), and α-COP-expression plasmid in pMX vector (200 ng) Twenty-four hours after the transfection, the luciferase activity in the total cell lysate was measured.

### Immunocytochemistry

Cells were fixed with 4% paraformaldehyde (PFA) in PBS at room temperature for 15 min, permeabilized with 0.1% Triton X-100 in PBS at room temperature for 5 min, and quenched with 50 mM NH_4_Cl in PBS at room temperature for 10 min. After blocking with 3% BSA in PBS, cells were incubated with primary antibodies, then with secondary antibodies conjugated with Alexa fluorophore.

### Confocal microscopy

Confocal microscopy was performed using a LSM880 with Airyscan (Zeiss) with a 63 x 1.4 Plan-Apochromat oil immersion lens.

### Immunoprecipitation

Cells were lysed with IP buffer (50 mM HEPES-NaOH (pH 7.2), 150 mM NaCl, 5 mM EDTA, 1% CHAPS, protease inhibitors (Protease Inhibitor Cocktail for Use with Mammalian Cell and Tissue Extracts, 25955-11, nacalai tesque), and phosphatase inhibitors (8 mM NaF, 12 mM beta-glycerophosphate, 1 mM Na_3_VO_4_, 1.2 mM Na_2_MoO_4_, 5 µM cantharidin, and 2 mM imidazole). The lysates were centrifugated at 15000 rpm for 10 min at 4 °C, the resultant supernatants were incubated for overnight at 4 °C with anti-FLAG M2 Affinity Gel or anti-GFP nanobody beads^26^ for 1 h. The beads were washed four times with immunoprecipitation wash buffer (50 mM HEPES-NaOH (pH 7.2), 150 mM NaCl, 0.7% CHAPS), and eluted with elution buffer (50 mM HEPES-NaOH (pH 7.2), 150 mM NaCl, 5 mM EDTA, 1% Triton X-100, 200 µg/mL FLAG peptide).

### Immunoelectron microscopy

Cells were fixed with 4% PFA (1.04005.1000, MERCK), 4% sucrose and 0.1 M phosphate buffer (pH 7.2) for 10 min at room temperature and then 30 min at 4 °C. After rinsing with 7.5% sucrose and 0.1 M phosphate buffer (pH 7.4), they were scraped and embedded in 10% gelatin (G2500, Sigma) and 0.1 M phosphate buffer (pH 7.4). The cell blocks were cut into small pieces (about 1 mm cube), which were infused overnight with 20% polyvinylpyrrolidone (PVP10, Sigma-Aldrich), 1.84 M sucrose, 10 mM Na_2_CO_3_ and 0.08 M phosphate buffer (pH 7.4) followed by rapid freezing in liquid nitrogen^27^. Ultrathin cryosections were prepared using an ultramicrotome (EM UC7, Leica) equipped with a cryochamber (EM FC7, Leica). They were incubated with 1% BSA and PBS for 20 min at room temperature, and then with the primary antibody against GFP (ab6556, dilution 1:100) (Abcam) for 24 h at 4 °C. After incubation with 12 nm colloidal gold particle-conjugated donkey anti-rabbit (711-205-152, dilution 1:20) (Jackson ImmunoResearch) for 1 h at room temperature, they were fixed with 2% glutaraldehyde (G017/1, TAAB) and PBS for 5 min. They were stained with 2% uranyl acetate for 5 min and embedded in 0.17% uranyl acetate and 0.33% polyvinyl alcohol (P8136, Sigma-Aldrich). After drying up, sections were observed using an electron microscope (JEM1200EX, JEOL).

### Western blotting

Proteins were separated in polyacrylamide gel and then transferred to polyvinylidene difluoride membranes (Millipore). These membranes were incubated with primary antibodies, followed by secondary antibodies conjugated to peroxidase. The proteins were visualized by enhanced chemiluminescence using a LAS-4000 (GE Healthcare).

### Mass spectrometry

Cells were lysed with IP buffer (50 mM HEPES-NaOH (pH 7.2), 150 mM NaCl, 5 mM EDTA, 1% Triton X-100, protease inhibitors, and phosphatase inhibitors). The lysates were centrifugated at 15000 rpm for 10 min at 4 °C, the resultant supernatants were incubated for overnight at 4 °C with anti-FLAG M2 Affinity Gel. The beads were washed four times with immunoprecipitation wash buffer (50 mM HEPES-NaOH (pH 7.2), 150 mM NaCl, 1% Triton X-100), and eluted with elution buffer (50 mM HEPES-NaOH (pH 7.2), 150 mM NaCl, 5 mM EDTA, 1% Triton X-100, 500 µg/mL FLAG peptide. Eluted proteins were applied to SDS-PAGE, and the electrophoresis was stopped when the samples were moved to the top of the separation gel. The gel was stained with CBB and the protein bands at the top of separation gel were excised. The proteins were reduced and S-carboxylmethylated, followed by a tryptic digestion in gel (TPCK treated trypsin, Worthington Biochemical Corporation). The digests were separated with a reversed phase nano-spray column (NTCC-360/75-3-105, NIKKYO technos) and then applied to Q Exactiv Hybrid Quadrupole-Orbitrap mass spectrometer (Thermo Scientific). MS and MS/MS data were obtained with TOP10 method. The MS/MS data was searched against NCBI nr database using MASCOT program 2.6 (Matrix Science) and the MS data was quantified using Proteome Discoverer 2.2 (Thermo Scientific).

### RNA interference

siRNA specific to Ablim1 (M-059643-01), Clptm1l (M-062856-00), Ddost (M-064791-00), H13 (M-059025-01), Hsd17b12 (M-060708-01), Lpcat1 (M-059984-01), Mars (M-066281-00), Myl12a (M-046975-01), Ncl (M-059054-01), Ncln (M-052038-01), Psmd11 (M-057766-01), Hacd3 (M-065373-01), Slc25a4 (M-061103-01), Slc25a5 (M-042392-01), Smpd4 (M-042323-01), Stt3b (M-062290-01), Surf4 (M-062783-00), Tmem43 (M-046911-01), were purchased from Dharmacon. siRNA specific to Surf4 (Surf4 Stealth Select RNAi) purchased from Thermo Fisher Scientific. Negative control siRNA was purchased from Dharmacon and Thermo Fisher Scientific. A total of 20 nM siRNA was introduced to cells using Lipofectamine RNAiMAX (Invitrogen) according to the manufacturer’s instruction. After 6 h, the medium was replaced by DMEM with 10% heat-inactivated FBS and cells were further incubated for 44 or 68 h for subsequent experiments.

### Statistical analyses

Error bars displayed throughout this study represent s.e.m. unless otherwise indicated, and were calculated from triplicate or quadruplicate samples. Statistical significance was determined with one-way ANOVA followed by Tukey– Kramer post hoc test.; \**P* < 0.05; \*\**P* < 0.01; \*\*\**P* < 0.001; NS not significant (P > 0.05). Data shown are representative of 2–3 independent experiments, including microscopy images and Western blots.

