## Supplementary Figures for "Homeostatic regulation of STING by Golgi-to-ER membrane traffic"

### Supplementary Figure 1

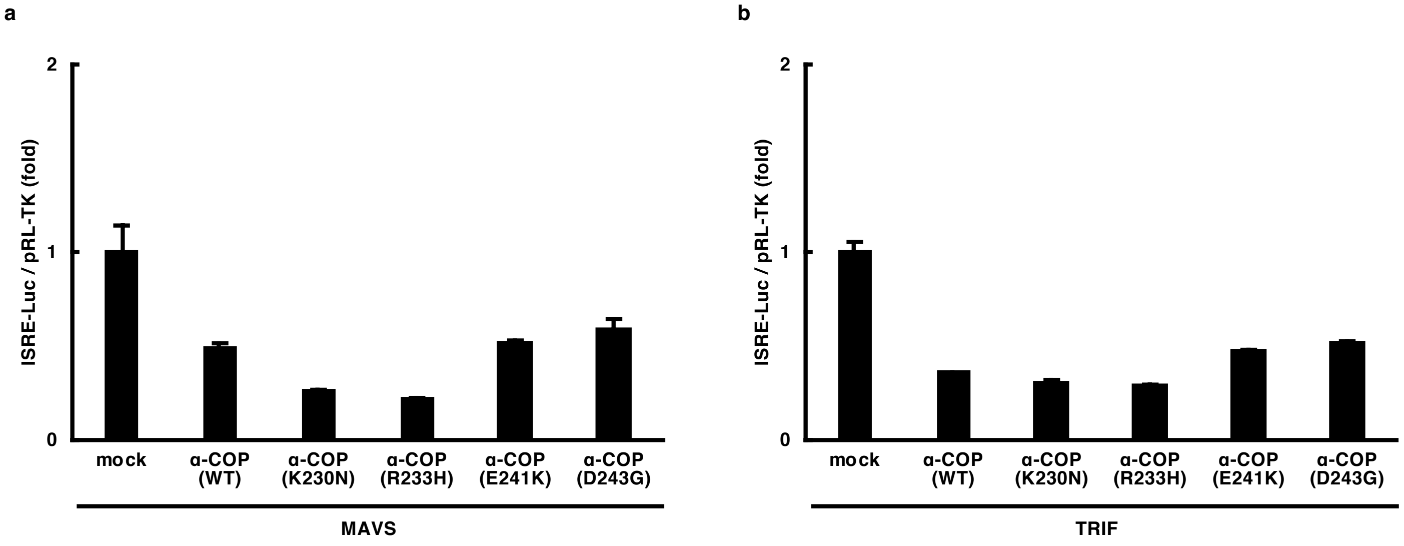


**Supplementary Figure 1 l The α-COP variants do not activate the IRF3 promoter in cells transfected with MAVS or TRIF.** (**a**,**b**) HEK293T cells were transfected as indicated, together with the ISRE (also known as PRDIII or IRF-E)-luciferase reporter. Luciferase activity was then measured. Data represent mean s.e.m. of three independent experiments.

### Supplementary Figure 2

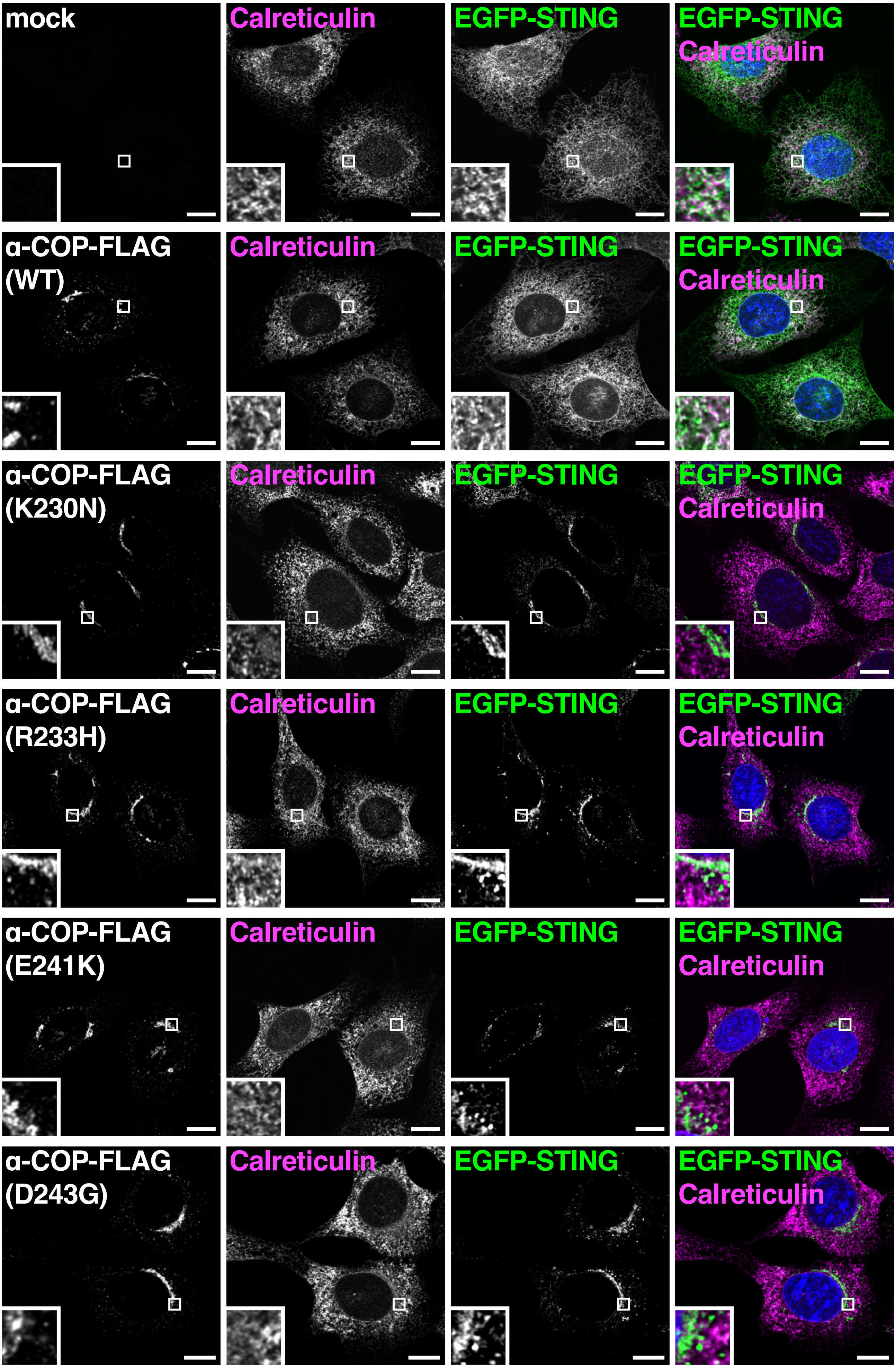


**Supplementary Figure 2 | STING translocates from the ER in the α-COP variants-expressing cells.** α-COP-FLAG and EGFP-STING were stably expressed in *Sting*$^{-/-}$ MEF cells. Cells were fixed, permeabilized, and stained for calreticulin (an ER protein). Nuclei were stained with DAPI (blue). Scale bars, 10 $\mu$m.

### Supplementary Figure 3

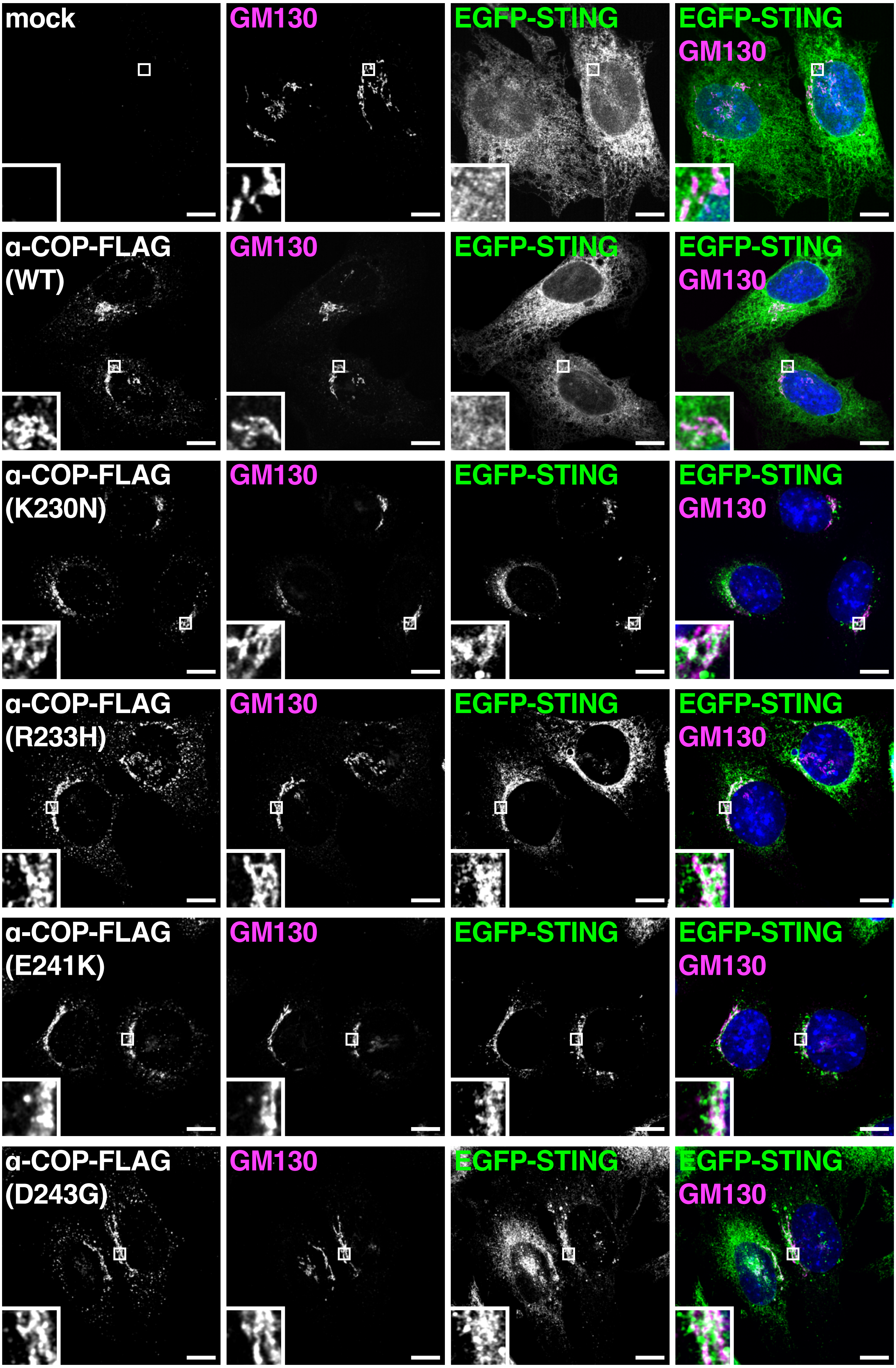


**Supplementary Figure 3 I STING translocates to the Golgi in the α-COP variants-expressing cells.** α-COP-FLAG and EGFP-STING were stably expressed in *Sting*$^{-/-}$ MEF cells. Cells were fixed, permeabilized, and stained for GM130 (a Golgi protein). Nuclei were stained with DAPI (blue). Scale bars, 10 $\mu$m.

### Supplementary Figure 4

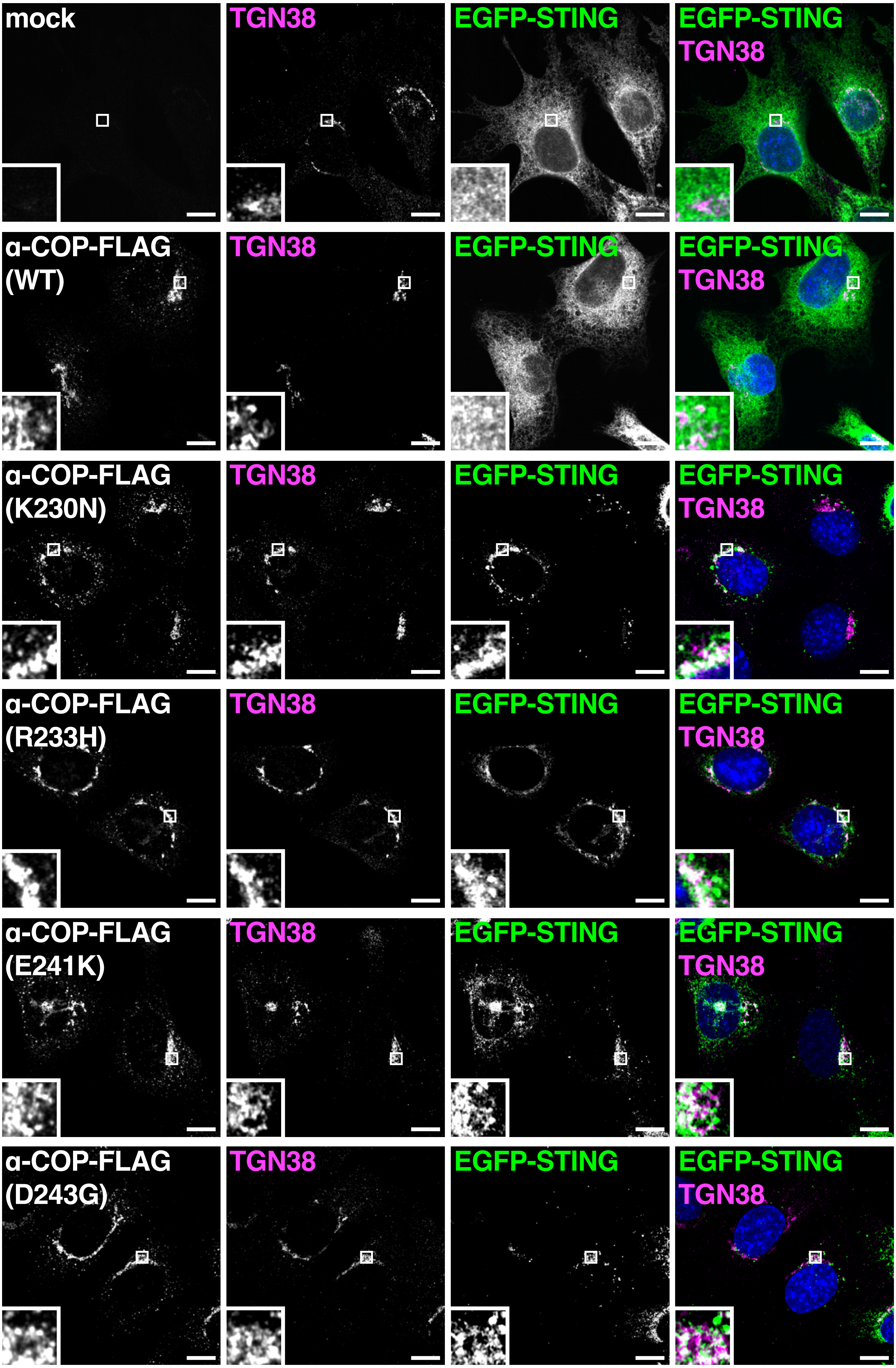


**Supplementary Figure 4 I STING translocates to the Golgi in the α-COP variants-expressing cells.** High resolution images of Fig. 2a.

### Supplementary Figure 5

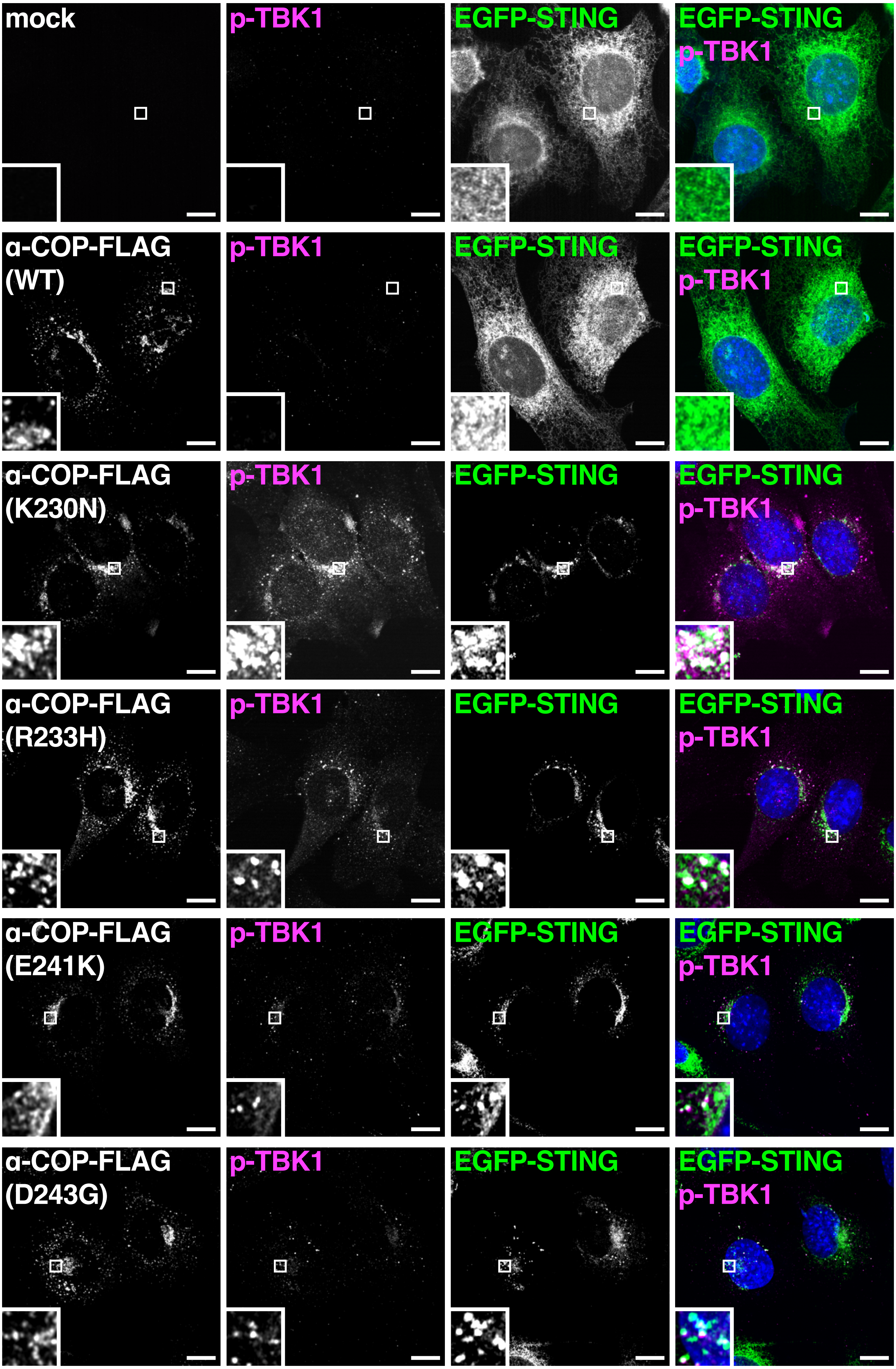


**Supplementary Figure 5 | STING activates TBK1 at perinuclear compartments in the α-COP variants-expressing cells.** High resolution images of Fig. 2b.

### Supplementary Figure 6

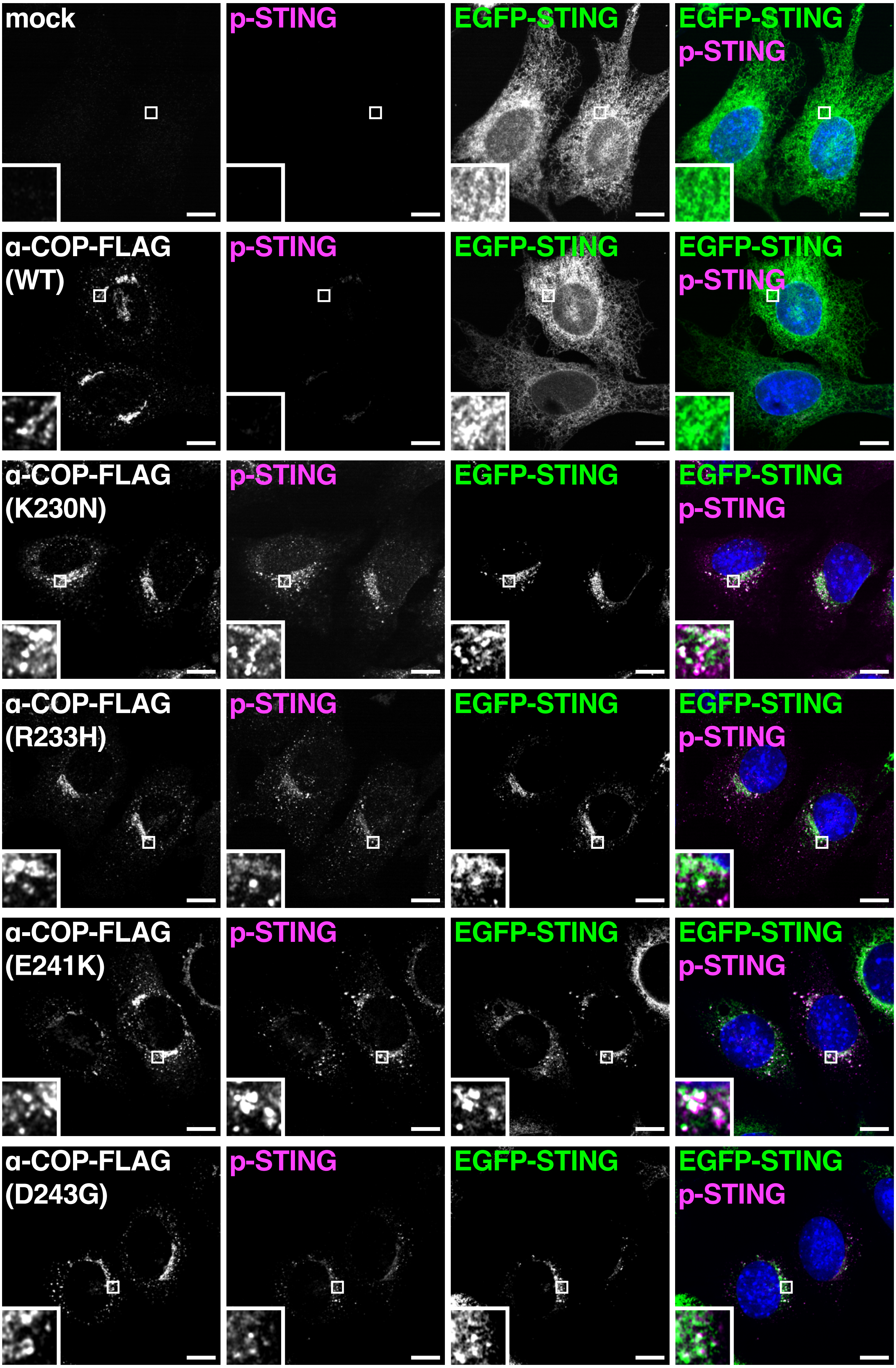


**Supplementary Figure 6 | Phosphorylated STING at Ser365 is detected at perinuclear compartments in the α-COP variants-expressing cells.** α-COP-FLAG and EGFP-STING were stably expressed in *Sting*$^{-/-}$ MEF cells. Cells were fixed, permeabilized, and stained for phospho-STING (p-Ser365). Nuclei were stained with DAPI (blue). Scale bars, 10 μm.

### Supplementary Figure 7

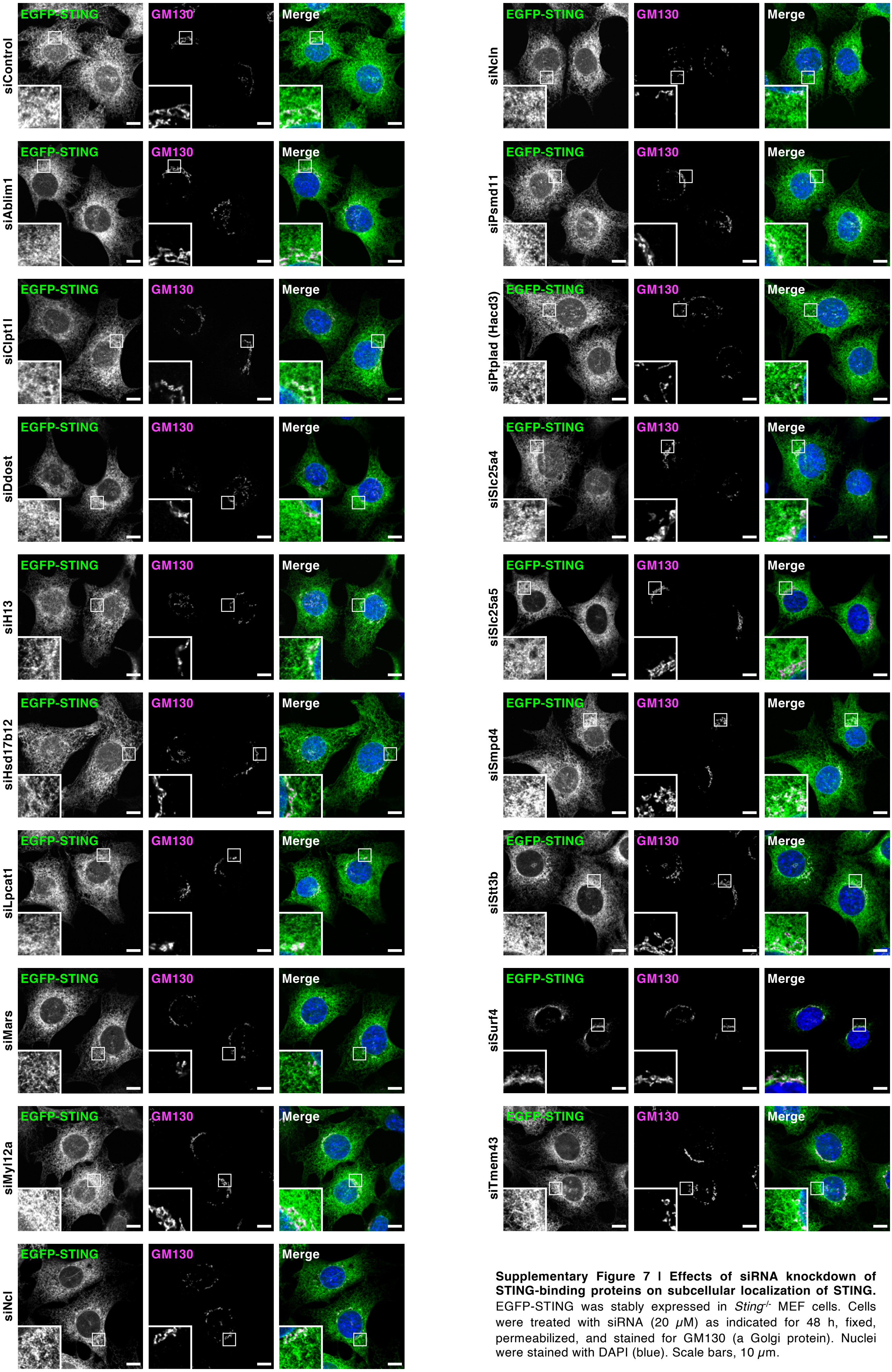


**Supplementary Figure 7 | Effects of siRNA knockdown of STING-binding proteins on subcellular localization of STING.** EGFP-STING was stably expressed in *Sting*$^{-/-}$ MEF cells. Cells were treated with siRNA (20 $\mu$M) as indicated for 48 h, fixed, permeabilized, and stained for GM130 (a Golgi protein). Nuclei were stained with DAPI (blue). Scale bars, 10 $\mu$m.

### Supplementary Figure 8

**a**

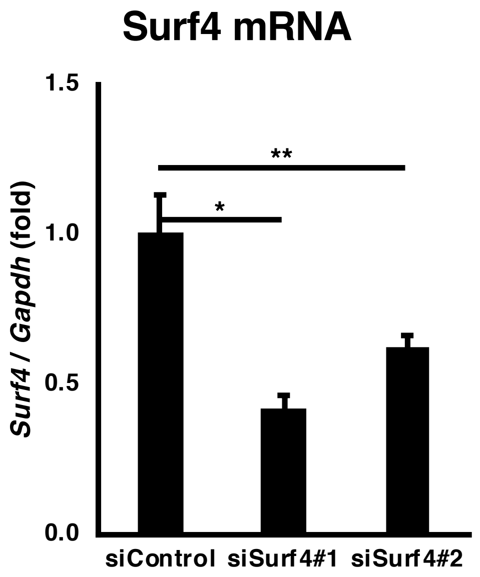


**b**

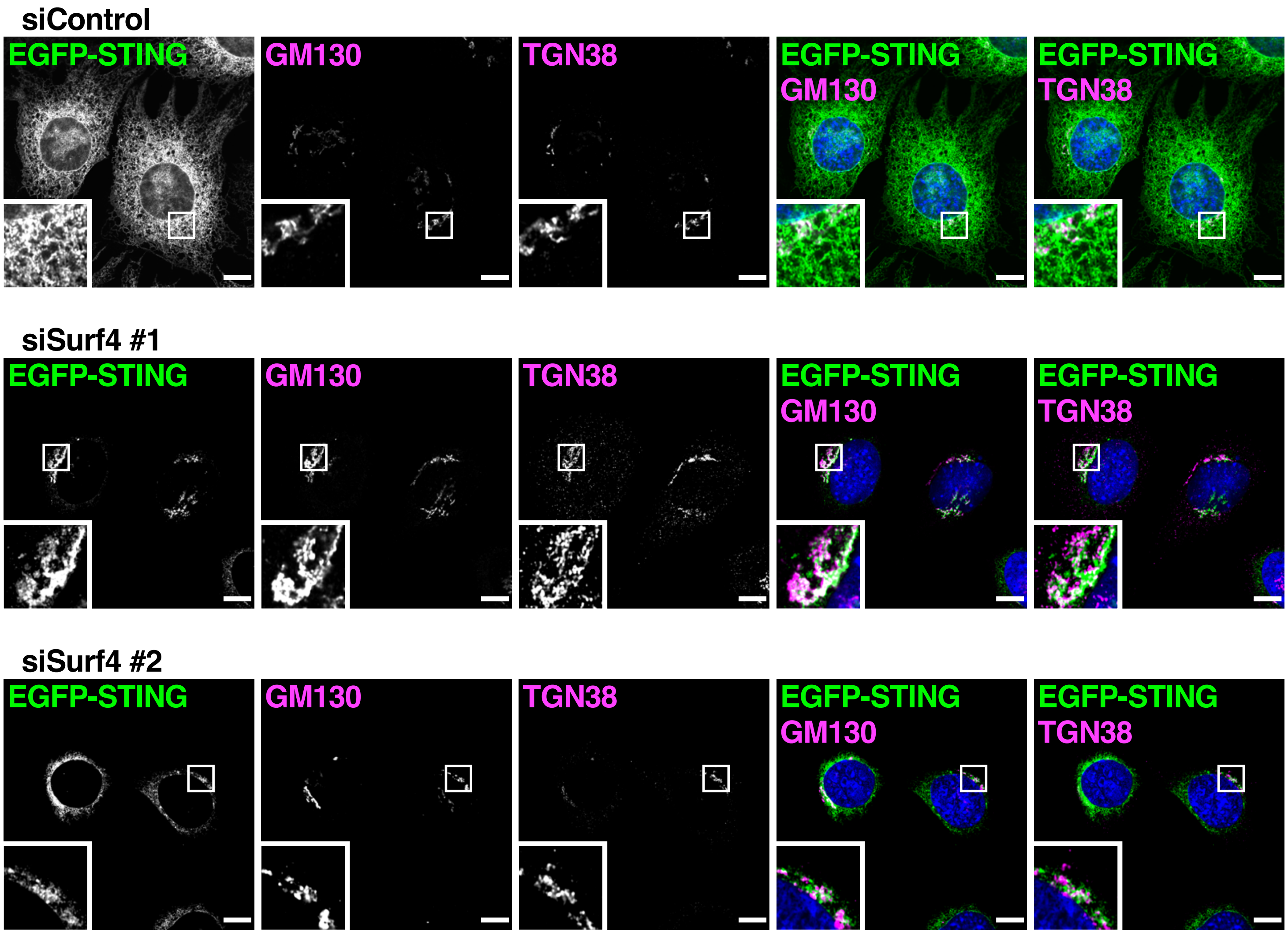


**Supplementary Figure 8 l Knockdown experiments of Surf4.** (**a**) EGFP-STING was stably expressed in *Sting*$^{-/-}$ MEF cells. Cells were treated with siRNA (20 $\mu$M) as indicated for 48 h and qRT-PCR of the expression of Surf4 was performed. Data represent mean s.e.m. of three independent experiments. **P*<0.01, ***P*<0.005 (one-way analysis of variance). (**b**) High resolution images of Fig. 3c.

### Supplementary Figure 9

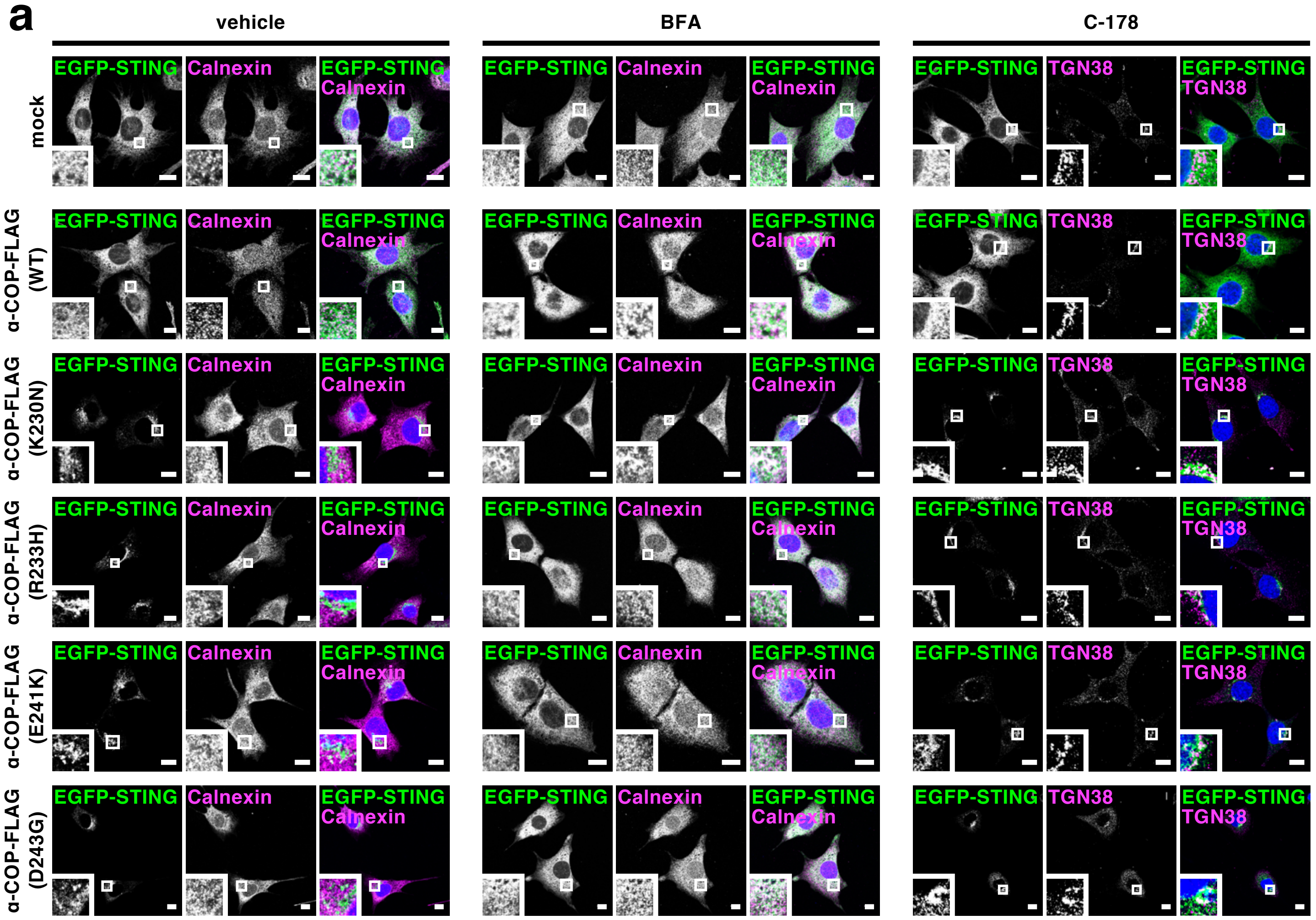


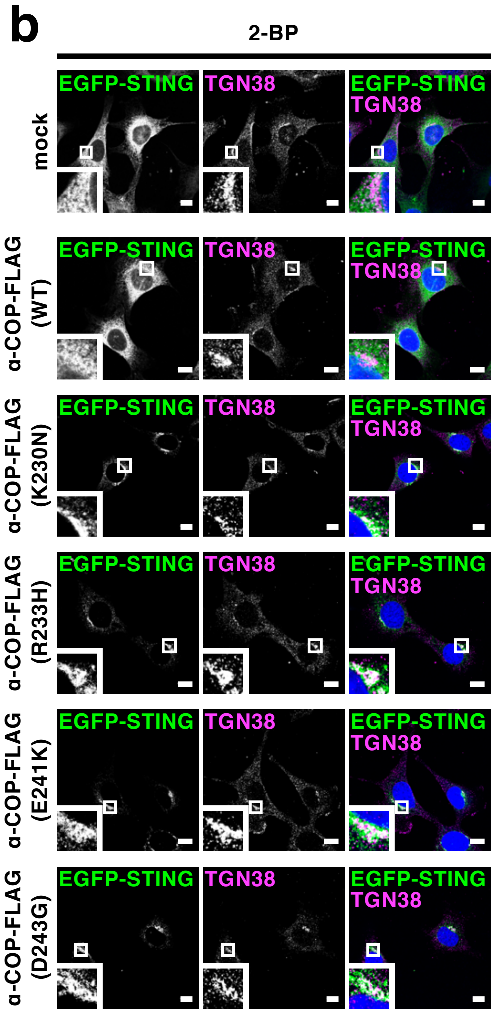


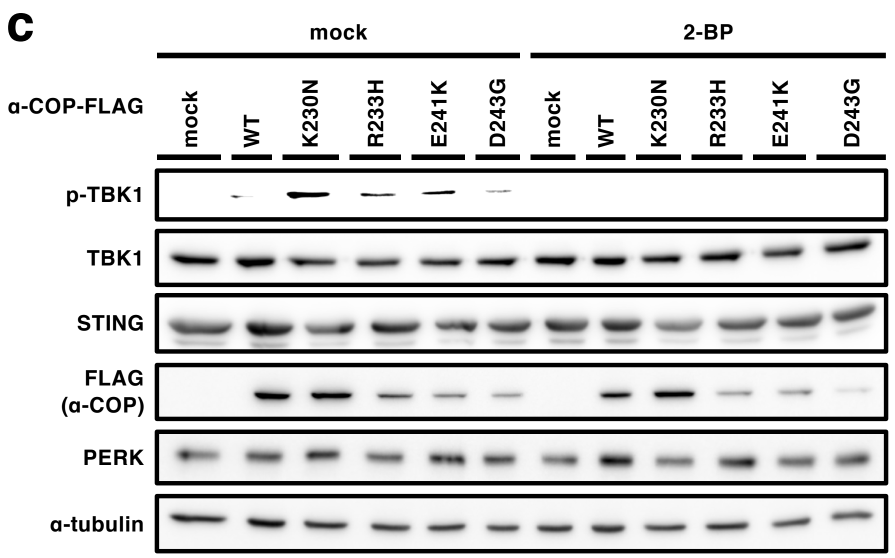


**Supplementary Figure 9 | The effect of BFA, C-178, or 2-BP treatment in the α-COP variants-expressing cells.** (**a**,**b**) α-COP-FLAG and EGFP-STING were stably expressed in *Sting*$^{-/-}$ MEF cells. Cells were treated with BFA (0.3 $\mu$g/mL), C178 (10 $\mu$M) (**a**), or 2-BP (100 $\mu$M) (**b**) for 8 h, fixed, permeabilized, and stained for calnexin (an ER protein) or TGN38 (a Golgi protein). Nuclei were stained with DAPI (blue). Scale bars, 10 $\mu$m. (**c**) α-COP-FLAG and EGFP-STING were stably expressed in *Sting*$^{-/-}$ MEF cells. Cells were treated with 2-BP (100 $\mu$M) for 8 h. Cell lysates were prepared and analyzed by western blot.

### Supplementary Figure 10

a

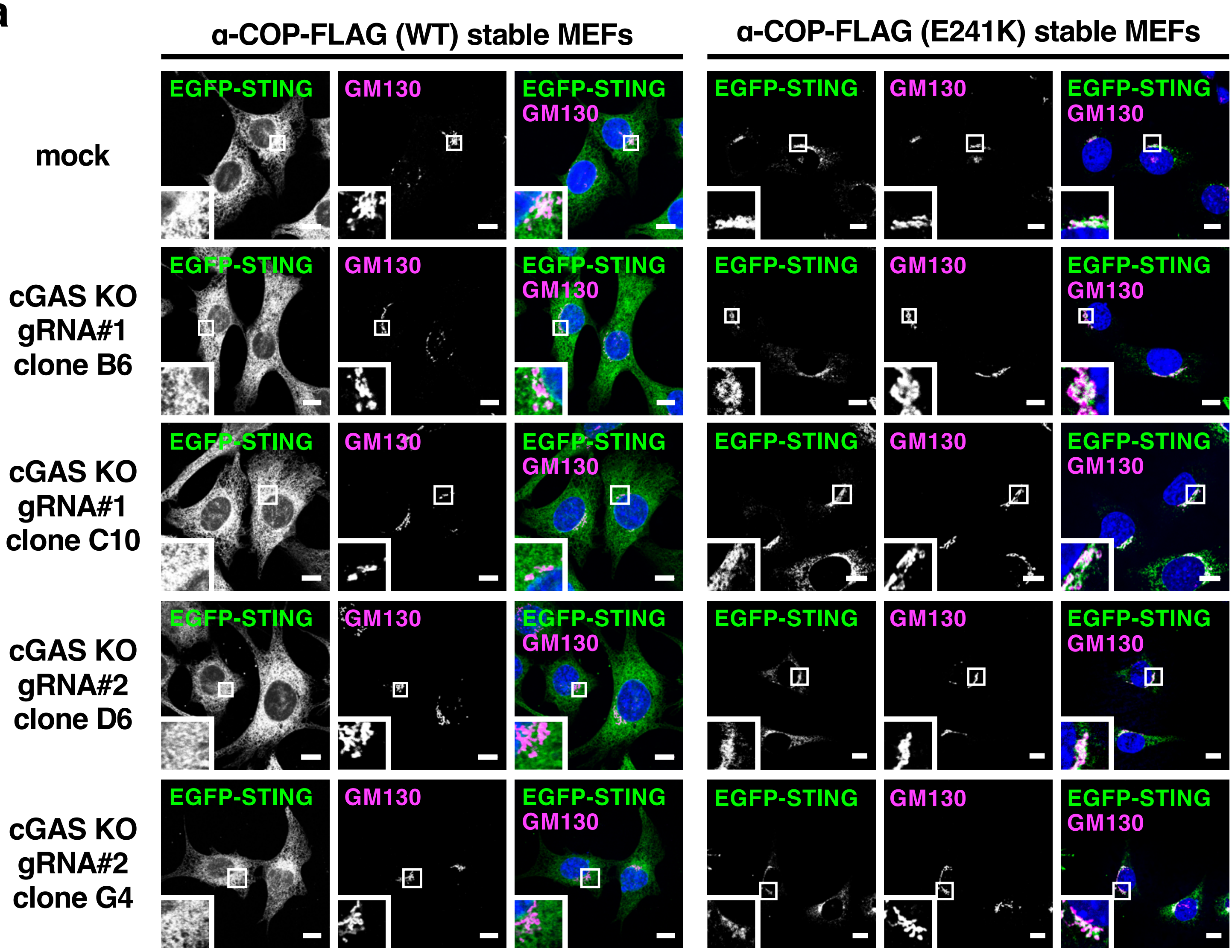


b

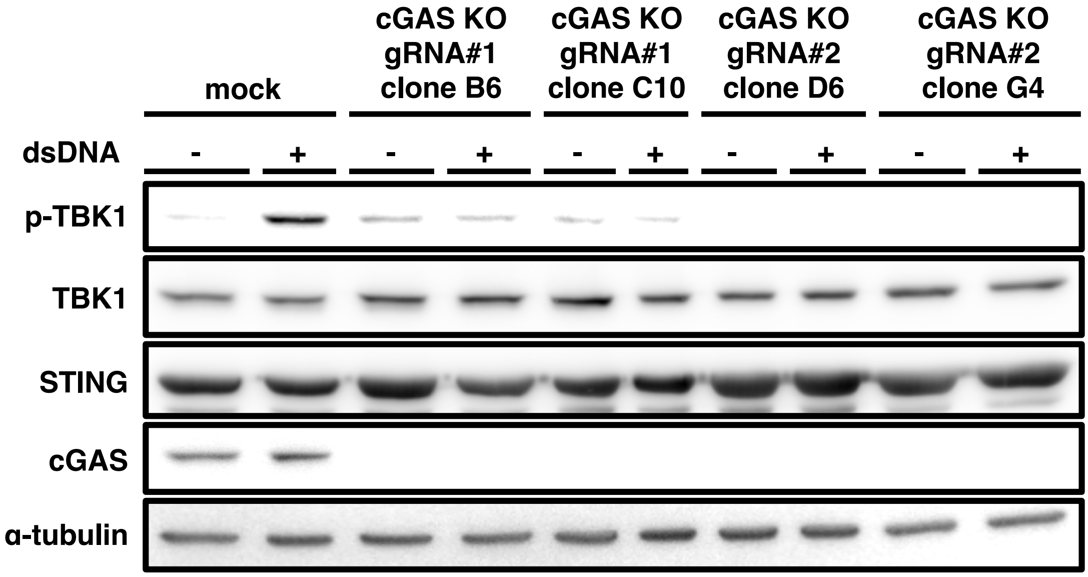


**Supplementary Figure 10 I STING translocates to the Golgi in the α-COP variants-expressing cGAS KO cells.** (**a**) cGAS knockout MEFs were generated by CRISPR-Cas9 system with *Sting*$^{-/-}$ MEFs. α-COP-FLAG and EGFP-STING were stably expressed in cGAS KO *Sting*$^{-/-}$ MEF cells. Cells were fixed, permeabilized, and stained for GM130 (a Golgi protein). Nuclei were stained with DAPI (blue). Scale bars, 10 $\mu$m. (**b**) Cells were stimulated with dsDNA for 2 h. Cell lysates were prepared and analyzed by western blot.
